# *Mammalian orthoreovirus* infection is enhanced in cells pre-treated with sodium arsenite

**DOI:** 10.1101/555367

**Authors:** Michael M. Lutz, Megan P. Worth, Meleana M. Hinchman, John S. L. Parker, EmilyD. Ledgerwood

## Abstract

Following reovirus infection, cells activate stress responses that repress canonical cellular translation as a mechanism to limit production of progeny virions. This includes the formation of stress granules (SG) that sequester translationally-stalled cellular transcripts, translation initiation factors, ribosomal proteins, and RNA binding proteins until conditions improve and translation can resume. Work by others suggests that these cellular stress responses, which are part of the integrated stress response, may benefit rather than repress reovirus replication. In agreement with this, we report that stressing cells prior to infection with sodium arsenite (SA), a robust inducer of SG and activator of eIF2α kinases, enhanced viral protein expression, percent infectivity and viral titer in SA-treated cells compared to untreated cells. SA-mediated enhancement of reovirus replication was not strain-specific, but was cell-type specific. While pre-treatment of cells with SA offered the greatest enhancement, treatment of infected cultures as late as 4 h post infection resulted in an increase in the percent of cells infected. SA activates the HRI kinase, which phosphorylates eIF2α and subsequently induces SG formation. Other stresses, such as heat shock (HS) and osmotic shock also activate HRI. Heat shock of cells prior to reovirus infection readily induced SG in greater than 85% of cells. Although HS pre-treatment had no effect on the percentage of infected cells or viral yield, it did enhance viral protein expression. These data suggest that SA pre-treatment perturbs the cell in a way that is beneficial for reovirus and that neither HRI activation nor SG induction is sufficient for reovirus infection enhancement.

**SIGNIFICANCE:** All viruses rely on the host translational machinery for the synthesis of viral proteins. In response to viral infection, cells activate the integrated stress response resulting in the phosphorylation of eIF2α and translation shutoff. Despite this, reovirus replicates to reduced titers in the absence of this response. In this work, we report that sodium arsenite activation of the integrated stress response prior to virus inoculation enhances virus infectivity, protein expression and titer. Together, these data suggest that modulation of conserved cellular stress responses can alter reovirus replication.

## INTRODUCTION

Acute viral infection induces stress within infected cells. The integrated stress response (ISR) is activated in many cells during viral infection [1-3]. The ISR facilitates cellular survival and a return to homeostasis, or initiates cell death signaling under conditions of severe stress or when the initiating stressor is maintained [4]. Four distinct stress kinases, can be activated in response to stress. Although, these kinases may have a number of substrates, they all phosphorylate the alpha subunit of the eukaryotic translation initiation factor 2 (eIF2α) at serine position 51. When eIF2α is phosphorylated, eIF2B is unable to exchange GDP for GTP, preventing eIF2 interaction with the Met-tRNAi and translation initiation [4].

The ISR is activated in response to accumulation of phosphorylated eIF2α. The four cellular kinases that phosphorylate eIF2α are heme-regulated eIF2α kinase (HRI), general control non-depressible 2 (GCN2), double-stranded RNA-dependent protein kinase (PKR) and PKR-like ER kinase (PERK) [4]. Both cell intrinsic and extrinsic stresses can activate these kinases: (i) HRI - heme-deprivation; (ii) GCN2 - amino acid starvation; (iii) PKR - accumulation of double-stranded RNA (as can occur during viral infection) and (iv) PERK - accumulation of unfolded proteins in the endoplasmic reticulum (ER) [4]. However, redundancy does exist amongst the kinases. For example, GCN2 is activated in response to ER stress in cells lacking PERK, and all four kinases can be activated under oxidative stress [5-9].

When the alpha subunit of eIF2 is phosphorylated at position 51, the GDP-loaded eIF2 entraps the limiting amounts of eIF2B leading to a rapid decrease in concentration of functional eIF2.GTP.Met-tRNAi ternary complexes. This results in reduced translation initiation and inhibition of de novo protein synthesis [4]. In response to the inhibition of translation initiation, stress granules (SG) form in the cytoplasm. SG assembly is mediated by the RNA binding proteins T-cell antigen 1 (TIA-1), TIA-1 related protein (TIAR) and the Ras-GAP SH3-binding protein 1 (G3BP1), and results in the compartmentalization of translationally-stalled mRNA transcripts, RNA-binding proteins, 40S ribosomes, and translation initiation factors [10, 11]. SG are considered to be sites of mRNA triage, protecting mRNA transcripts until a stress is alleviated and the cell returns to homeostasis.

Many viruses prevent activation of the integrated stress response to maintain protein translation and to ensure successful viral infection. To do this, viruses target the initiating kinases and/or the downstream effector mechanisms of the ISR. Cells in which the initiating kinases, such as PKR are inhibited or knocked out are often more permissive for viral replication [12, 13]. However, not all viruses benefit from inhibition of kinase activity and initiation of the ISR. A study examining the role of the PKR-eIF2 pathway during mammalian orthoreovirus (reovirus) infection found some strains had reduced titers in PKR knockout murine embryonic fibroblasts (MEF) [14]. Follow-up studies observed both increased ISR gene expression and reduced levels of the eIF2α kinase inhibitor P58^IPK^ in cells infected with reovirus strains known to robustly interfere with host translation, and these strains replicated less efficiently in MEFs expressing a non-phosphorylatable form of eIF2α [15].

Reovirus infection also modulates stress granule formation that occurs downstream of ISR activation [15, 16]. Early in infection, entering viral core particles localize to stress granules that form within infected cells. However, within 4-6 h after infection, the stress granules have disappeared and viral factories (VFs), the sites of reovirus replication, transcription, translation and assembly, become prominent [16]. In some reovirus-infected cells, the stress granule protein G3BP1 localizes to the margins of the VFs, mediated by an interaction of G3BP1 with the non-structural viral protein σNS [17]; σNS interacts with the nonstructural protein µNS that forms the matrix of VFs [18]. Co-expression of σNS and μNS is sufficient to alter the localization of G3BP1 and suppress stress granule induction [17]. The interplay between eIF2α phosphorylation, PKR activation, translational shutoff and G3BP1 induced SG formation is strain-dependent, with SG formation negatively impacting some strains of reovirus [17]. Together, these studies suggest a unique role for the ISR during reovirus infection, however the magnitude of this role remains to be elucidated. Most virological studies of the ISR take two approaches: 1) understanding the impact of virus infection on stress responses or 2) understanding how perturbing the ISR during infection affects the virus. Given the previous observation that reovirus replicates to lower titers in cells with an impaired ISR, we hypothesized that reovirus infection would be enhanced in cells in which the ISR has been activated prior to infection. We found that reovirus infection was more efficient (increased infectivity, protein expression, and replication) in cells in which the ISR had been activated by pre-treatment with sodium arsenite prior to virus adsorption. Sodium arsenite-induced enhancement of reovirus infection was observed in all reovirus strains tested but was dependent on cell-type and the time of sodium-arsenite addition. Enhancement of viral infectivity was only observed if sodium arsenite was added to cells within 4 h of inoculation, with maximal enhancement if the addition occurred prior to inoculation, suggesting a relationship between the ISR and early replication events. Furthermore, not all activators of the ISR were equally beneficial as heat shock prior to infection had no impact on viral replication. Taken together, these data suggest a critical role for the ISR during reovirus infection and that activation of the ISR prior to reovirus infection is beneficial in some cell-types.

## MATERIALS AND METHODS

### Cells and reagents

CV-1 (ATCC CCL-70) and HeLa cells were maintained in Eagles minimum essential medium (MEM) (CellGro) containing 10% fetal bovine serum (FBS; Hyclone), 100 mM sodium pyruvate (CellGro), and 200 mM L-glutamine (CellGro) at 37°C in the presence of 5% CO_2_. L929 cells were maintained in MEM containing 8% FBS and 200 mM L-glutamine at 37°C in the presence of 5% CO_2_. Human Pancreatic Ductal Epithelial (HPDE) cells (Kerafast H6c7) were maintained in keratinocyte SFM (serum-free medium) supplemented with 25 mg bovine pituitary extract and 2.5 μg human recombinant epidermal growth factor, both provided (Invitrogen) at 37°C in the presence of 5% CO_2_. Sodium arsenite was kindly donated by Dr. Shu-bing Qian (Cornell University) and was used at a final concentration of 0.5 mM in all experiments.

### Viruses

Reoviruses T1L and T3D laboratory stocks originated from the T1/human/Ohio/Lang/1952 and T3/human/Ohio/Dearing/1955 isolates respectively [19]. The superscript T3D^N^ differentiates a clone obtained from M.L. Nibert (Harvard Medical School) from T3D^C^ a clone obtained from L.W. Cashdollar (Medical College of Wisconsin). The two clones have been shown to differ in M1 gene sequence and viral factory morphology. All infections were performed with T3D^N^ (abbreviated T3D). The prototype reovirus serotype 3 strain Abney (T3A) was a kind gift from Dr. Barbary Sherry (North Carolina State University) and was plaque-purified and passaged twice on L-cell monolayers to generate working stocks.

### Infections

CV-1, L929, HeLa or HPDE cells were seeded in 24-well culture plates containing 12 mm glass coverslips or 12-well culture plates the day before to give rise to 50 to 80% confluence prior to infection. Cells were infected with virus at the indicated MOI for 1 h at room temperature (RT) in phosphate-buffered saline (PBS; pH 7.4), supplemented with 2 mM MgCl_2_, with rocking every 10 min. Following absorption, virus was removed and cells were incubated in growth medium at 37°C and harvested at the indicated time points.

### Antibodies

The following commercial primary antibodies were used for immunoblotting: mouse monoclonal anti-G3BP1 (2F3) antibody (H00010146-M01; Novus Biologicals), mouse monoclonal anti-TIAR antibody (sc-398372; Santa Cruz Biotechnology), and mouse monoclonal anti-alpha-tubulin antibody (NB100-690; Novus Biologicals). Reovirus protein expression was assessed using a chicken polyclonal antiserum against μNS prepared against bacterially-expressed purified antigen by Covance. Secondary antibodies used for immunoblotting were as follows: HRP Donkey Anti-Mouse IgG (715-035-150; Jackson ImmunoResearch), and HRP Donkey Anti-Chicken IgY (703-035-155; Jackson ImmunoResearch). Primary antibodies used for immunofluorescence included: mouse monoclonal anti-G3BP1 (611126; BD Biosciences), rabbit monoclonal anti-TIAR (8509S; Cell Signaling Technology), and chicken polyclonal antiserum against µNS to detect viral factories. Secondary Antibodies used for immunofluorescence assays included Alexa Fluor 594 goat anti-chicken IgG, Alexa Fluor 594 goat anti-rabbit IgG, Alexa Fluor 488 goat anti-mouse, and Alexa Fluor 488 goat anti-rabbit (Thermofisher).

### Stress granule induction

Stress granules were induced in cells using the following mechanisms: 1) treatment with 0.5 mM sodium arsenite in normal growth media at 37°C for 30 min prior to infection or 30 min prior to harvesting or 2) heat shock in normal growth media in a pre-heated incubator at 44 °C for 45 min prior to infection or harvesting. Cells were manually determined to contain stress granules if immunofluorescence revealed a minimum of three granules co-staining for both TIAR and G3BP.

### Immunofluorescence

Cells were washed once with PBS supplemented with 2 mM MgCl_2_ and fixed at room temperature for 10 min with 4% paraformaldehyde in PBS. Fixed cells were washed three times with PBS, permeabilized in 0.1% Triton X-100 in PBS for 15 min and washed three times with PBS. Cells were blocked for 15 min in staining buffer (SB; 0.05% saponin, 10mM glycine, 5% FBS, and PBS) and incubated with primary antibodies diluted in SB for 1 h. Cells were then washed one time with PBS before incubation with secondary antibodies diluted in SB for 1 h. Coverslips were mounted onto glass slides with ProLong Gold Anti-Fade reagent with DAPI (4[prime],6-diamidino-2-phenylindole; ThermoFisher). Images were obtained using an Olympus BX60 inverted microscope equipped with phase and fluorescence optics. Images were collected digitally with an Olympus DP74 color CMOS camera and cellSens Standard Software (Olympus) and were processed and prepared for presentation using photoshop (CC; Adobe) software.

### Immunoblot Assay

Cells were lysed in PBS containing 0.5% NP40, 140 mM NaCl, 30 mM Tris-HCl (pH 7.4), and an EDTA-free protease inhibitor cocktail (04693159001; Sigma Aldrich) for 30 min on ice. Cell lysates were resolved by SDS-PAGE and proteins were detected using antibodies as described above. Images were collected using a C-Digit digital scanner and Image Studio Digits software (Version 4, LiCor). When appropriate, immunoblots were incubated in stripping buffer (200 mM glycine, 3.5 mM SDS, 1% Tween 20, [pH 2.2]) and re-probed.

### Plaque Assay

Cells were infected as above before washing with PBS supplemented with 2 mM MgCl_2_ and incubated in growth medium at 37°C for the indicated times. Cells were subjected to two freeze/thaw cycles prior to the determination of viral titer by the standard reovirus plaque assay in L929 cells [20]. To determine the viral titer in PFU/ml, the following equation was used: PFU/ml = number of plaques / (D × V) where D = dilution factor and V = volume of diluted virus / well.

## RESULTS

### Infection with reovirus induces stress granule formation

Previous reports have suggested that stress granules form following infection with reovirus, but reports are conflicting as to the timing of stress granule presence in infected cells [15, 16]. To evaluate this further, we first confirmed that stress granules could form in CV-1 cells by treating uninfected cells with 0.5 mM sodium arsenite for 30 min. We then fixed and co-immunostained for TIAR and G3BP, two RNA binding proteins required for stress granule formation. We saw stress granules in ∼91% of treated CV-1 cells, confirming that these cells can form stress granules (Figure 1A, upper panels). We next infected CV-1 cells with reovirus at a multiplicity of infection (MOI) of 10 and fixed and immunostained cells at the indicated times post-infection (p.i.) for the presence of stress granules (TIAR), and viral factories (µNS) (Figure 1A, lower panels). Stress granules were absent in mock-infected cells and began appearing in infected cells around 2 h p.i. We found that stress granule formation in reovirus-infected CV-1 cells peaked around 6 h p.i. with 6.6% of cells containing stress granules. By 8 h p.i. the percentage of infected cells containing stress granules had dropped to 3.1% and by 18 h p.i., the level was no different from mock-infected cells (Figures 1A-B). A previous report showed that stress granules were present in reovirus-infected human prostate carcinoma DU145 cells at 19.5 h post infection [15]. Although we did not see stress granules in CV-1 cells at late times post-infection with reovirus at an MOI = 1, we found that G3BP and TIAR co-localized at the periphery of viral factories at 24 h p.i. when cells were treated with 0.5 mM sodium arsenite for 30 min immediately prior to harvest (Figure 1C). We occasionally saw a similar phenotype at 24 h p.i. in infected cells not treated with sodium arsenite (Figure 1C). These findings are consistent with those of Choudhury et al. who found that G3BP localizes to the viral factory periphery during reovirus infection [17]. While reovirus appears to modulate the localization of stress granule proteins at late times post infection, we observed only a modest increase in the expression level of stress granule protein expression in infected cells as compared to mock that was independent of MOI (Figure 1D). Together, these data suggest that reovirus infection induces stress granules at early, but not late times post-infection in CV-1 cells.

**Figure 1.**
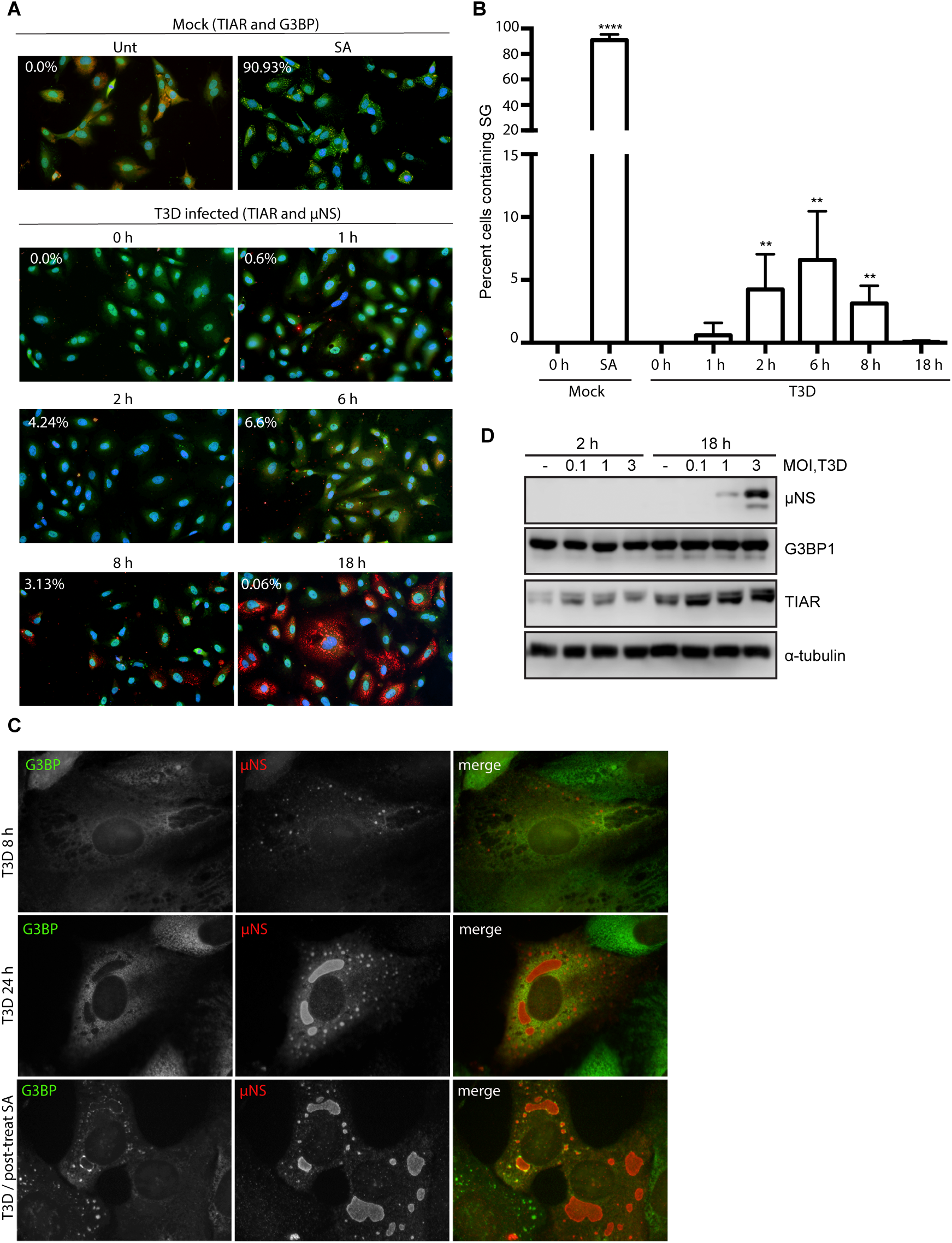
Reovirus infection induces the formation of stress granules at early times post infection. (A) CV-1 cells were mock infected, pre-treated with 0.5 mM sodium arsenite (SA) for 30 min, or infected with T3D at an MOI = 10. At the indicated times, cells were fixed and co-immunostained for G3BP (red) and TIAR (green), upper panels, to detect stress granules (SG) in uninfected cells or TIAR (green) and μNS (red), lower panels, to detect SG in T3D-infected cells. Cell nuclei were stained with DAPI (blue). Mock and SA-treated cells were fixed at 0 h. (B) The percent cells containing SG from panel A was quantified [(# of cells containing SG / total # of cells) × 100] from a minimum of three independent experiments. ** *P* < 0.01, **** *P* < 0.0001; two-tailed unpaired t test. (C) CV-1 cells were infected with T3D at an MOI = 0.5 and at 8 and 24 h p.i. cells were fixed and co-immunostained with G3BP (green) and μNS (red), upper panels. Alternatively, CV-1 cells were infected with T3D at an MOI = 1 and at 23.5 h cells were post-treated with 0.5 mM SA for 30 min. At 24 h p.i. cells were fixed and co-immunostained with G3BP (green) and μNS (red), lower panels. (D) Mock infected (-) or T3D infected cells were lysed at 2 or 18 h p.i and the expression level of the indicated proteins was determined by immunoblotting.

### Pre-treatment of cells with 0.5 mM sodium arsenite enhances reovirus infectivity

Following infection with reovirus, the viral protein μNS orchestrates the formation of viral factories, the sites of virus replication, assembly, and translation [21-23]. To facilitate translation, cellular factors including eIF3, eIF4G and ribosomal subunits are recruited to the viral factory [23]. Many of these initiation factors are similarly compartmentalized within stress granules [24]. To date, most studies have focused on the induction of stress granules in response to viral infection, or viral suppression of stress granule formation during an infection. To our knowledge, no studies have been performed to assess the impact of the presence of stress granules on reovirus infection. To explore this, we first examined the effect of stress granule presence on viral protein expression. CV-1 cells were either left untreated (-), or were treated with 0.5 mM sodium arsenite for 30 min before infection at an MOI = 1 (pre) or 30 min before harvest (post). Cell lysates were collected at 0, 6, 10, 18 and 24 h p.i. and the expression level of the non-structural viral protein μNS was determined (µNS was first detectable by 10 h p.i.). At 10, 18 and 24 h p.i., the expression levels of µNS were consistently higher in cells pre-treated with sodium arsenite when compared to untreated cells or cells treated with sodium arsenite 30 min before harvest (Figure 2A).

**Figure 2.**
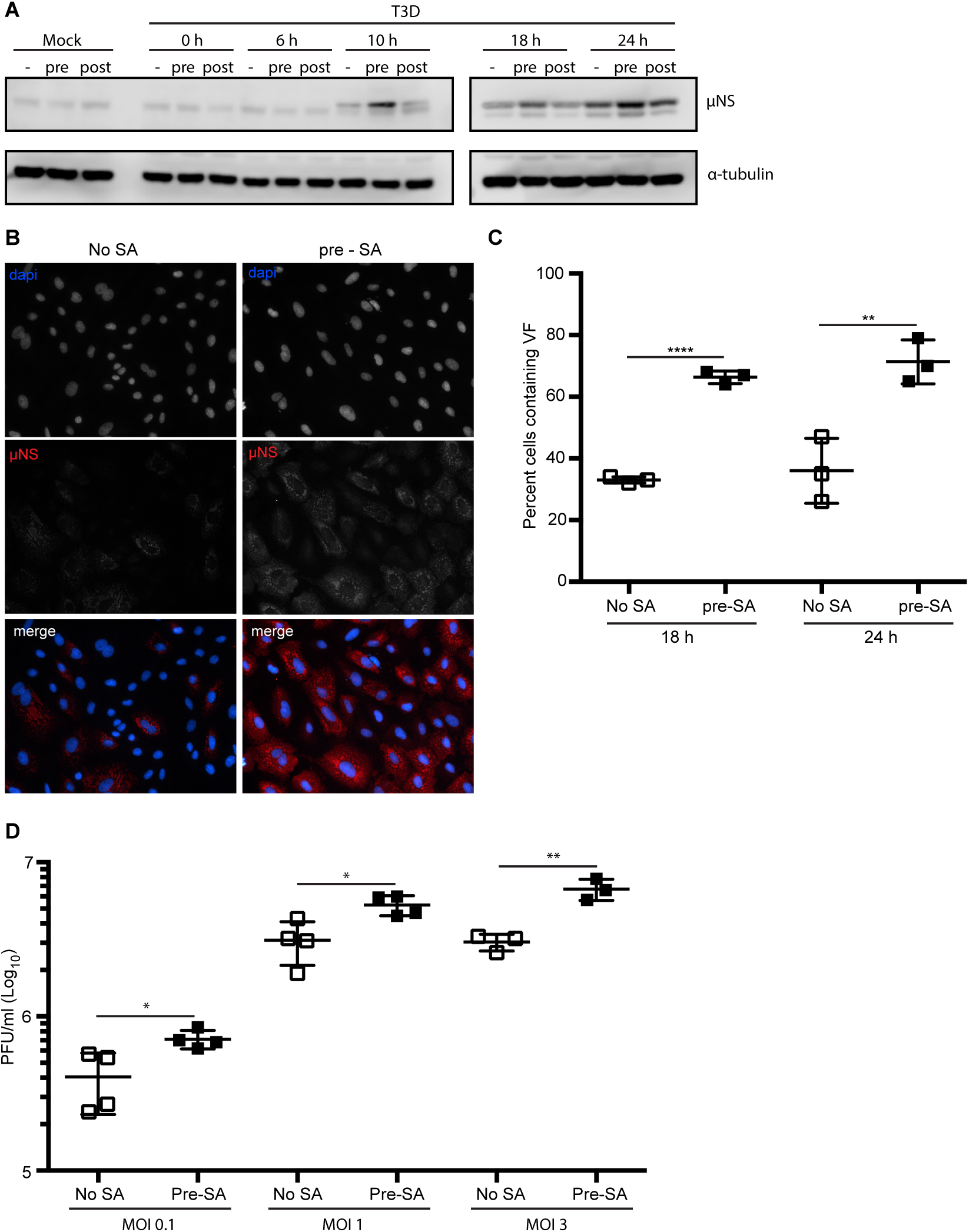
Pre-treatment with 0.5 mM sodium arsenite enhances reovirus infectivity. (A) CV-1 cells were mock infected or infected with T3D at an MOI = 1. Cells were left untreated (-), treated with 0.5 mM SA for 30 min prior to infection (pre) or were treated with 0.5 mM SA for 30 min immediately before lysis (post). At the indicated time points, cells were harvested and protein expression was determined by immunoblot. (B) CV-1 cells were infected with T3D at an MOI = 1. Cells were left untreated or treated with 0.5 mM SA for 30 min prior to infection (pre). At 18 h p.i., cells were fixed and immunostained with for µNS (red) and DAPI (nuclei, blue). (C) The percent cells containing viral factories (VF) as represented in panel B was quantified [(# of cells containing VF / total # of cells) × 100] from three independent experiments. (D) CV-1 cells were left untreated or were pre-treated with 0.5 mM SA for 30 min prior to infection with T3D at the indicated an MOI. At 18 h p.i., cells were subjected to the standard MRV plaque assay in L929 cells and plaques were counted from at least three independent experiments. * *P* < 0.05, ** *P* < 0.01, **** *P* < 0.0001; two-tailed unpaired t test.

As the viral protein μNS is necessary for formation of viral factories during reovirus infection, we next explored if the elevated μNS expression was a consequence of increased size of viral factories or increased numbers of infected cells [25]. We found that in cells pre-treated with 0.5 mM sodium arsenite for 30 min prior to reovirus infection, 67% of cells were infected (contained viral factories) at 18 h p.i., whereas only 33% of untreated cells were infected at 18 h p.i. (Figures 2B and C). We found similar findings at 24 h p.i. (Figure 2C).

Given the increased percentage of infected cells in dishes pre-treated with sodium arsenite, we tested if pre-treatment of cells with 0.5 mM sodium arsenite for 30 min before reovirus infection enhanced viral yield. We found that pre-treatment with sodium arsenite led to a modest but significant 2-3 fold increase in viral titer (PFU/ml) that was independent of multiplicity of infection, but consistent with the increased numbers of infected cells and the increased protein expression (Figure 2D). Together, these data indicate that pre-treatment of cells with sodium arsenite enhances reovirus infectivity by increasing the numbers of virus-permissive cells.

### SA-induced enhancement of reovirus infection is cell-type specific

Previous reports have suggested that reovirus-induced stress granule formation is cell-type specific [15, 16]. Therefore, we next sought to determine if the replication enhancement observed in CV-1 cells was seen in other cell types. We assessed the effect of pre-treatment with 0.5 mM sodium arsenite on CV-1, HeLa, L929 murine fibroblasts, and human pancreatic ductal epithelial (HPDE) cells on viral infectivity. In each cell line, a relative multiplicity of infection was chosen such that 20-50% of cells were infected. Consistent with our observations in CV-1 cells, pre-treatment with sodium arsenite in L929 and HPDE cells resulted in a higher percentage of cells exhibiting viral factories than untreated cells (Figure 3A). However, sodium arsenite pre-treatment did not increase the number of infected HeLa cells (Figure 3A). These data indicate that activation of the stress response pathways prior to infection through pre-treatment with sodium arsenite is beneficial in some, but not all, cell types.

**Figure 3.**
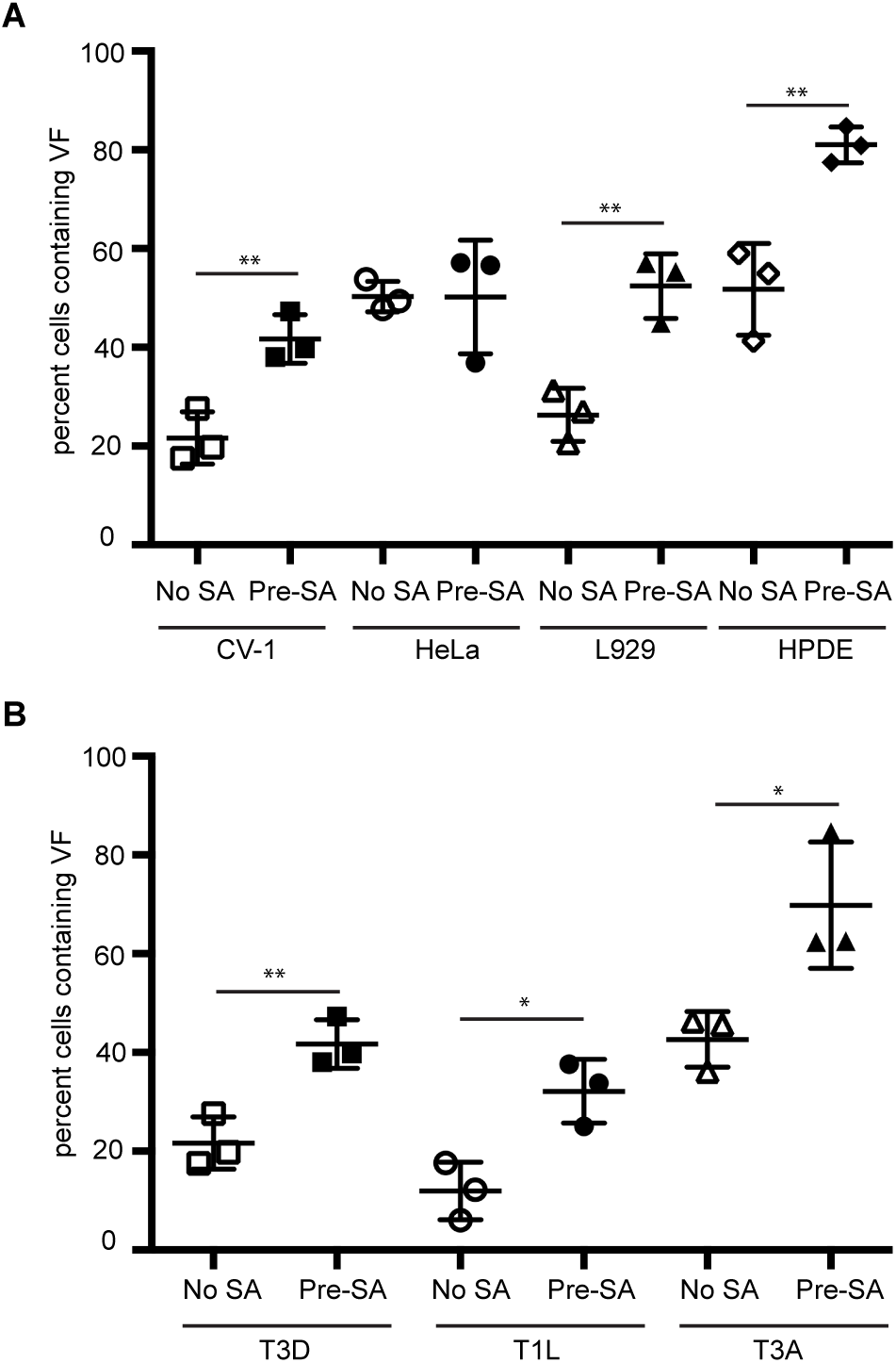
Pre-treatment with 0.5 mM sodium arsenite enhances permissivity in a cell-type-specific manner across reovirus strains. (A) CV-1, HeLa, L929 or HPDE cells were left untreated (No SA) or were treated with 0.5 mM SA for 30 min prior to infection (Pre-SA). Following this, cells were infected with T3D at an MOI = 1. At 18 h p.i., cells were fixed and immunostained for µNS (red) and DAPI (nuclei, blue). The percent of cells containing viral factories (VF) was quantified [(# of cells containing VF / total # of cells) × 100] from three independent experiments. (B) CV-1 cells were left untreated (No SA) or were treated with 0.5 mM SA prior to infection (Pre-SA). Cells were then infected with reovirus strains T3D, T1L, or T3A such that ∼20% of cells were infected. At 18 h p.i., cells were fixed and immunostained for µNS (red) and DAPI (nuclei, blue). The percent of cells containing VF was quantified [(# of cells containing VF / total # of cells) × 100] from three independent experiments. * *P* < 0.05; ** *P* < 0.01; two-tailed unpaired t test.

### SA-induced enhancement of reovirus infection is strain-independent

The prototypic reovirus strains Type 1 Lang (T1L), Type 2 Jones (T2J), Type 3 Abney (T3A), and Type 3 Dearing (T3D) differ in their capacity to induce host translational shutoff and to induce eIF2α phosphorylation [15, 26]. To determine if the benefit of pre-treatment with sodium arsenite was reovirus strain specific, we infected CV-1 cells with T3D, T1L or T3A and assessed the percentage of infected cells at 18 h p.i. compared to untreated cells. As observed following infection with the T3D strain, pre-treatment with sodium arsenite resulted in nearly a 2-fold increase in the percentage of T1L-and T3A-infected cells (Figure 3B). Sodium arsenite pre-treatment in L929 cells infected with T3D, T1L or T3A yielded similar results, however sodium arsenite pre-treatment had no effect on any strain tested in HeLa cells (Supplemental Figure 1). These data suggest that reovirus strains T3D, T3A and T1L benefit from pre-treatment with sodium arsenite prior to infection in some cell types.

### Heat shock prior to infection enhances viral protein expression in T3D-infected L-cells, but does not affect the percentage of infected cells or viral yield

Of the four stress-responsive kinases, sodium arsenite treatment activates HRI kinase. HRI is activated in erythrocytes by reduced heme availability, but it is ubiquitously expressed in many tissues and, when activated by sodium arsenite, phosphorylates eIF2α and subsequently induces stress granule formation [27]. HRI is the only stress kinase required for translational inhibition in response to arsenite treatment in mouse embryonic fibroblasts [27]. HRI kinase can also be activated by other cellular stresses including osmotic stress and heat shock. To further assess if the enhanced reovirus infectivity following sodium arsenite treatment was a consequence of HRI activation, we assessed viral protein expression, the percentage of infected cells and viral yield in response to heat shock. We used stress granule induction in response to heat shock as an indirect measure of HRI activation (Figure 4A). Our attempts to induce stress granules in CV-1 cells with heat shock were only able to achieve no greater than 25% of cells containing stress granules. Because of this, we assessed the effects of heat shock induction of stress granules in L929 cells. Heat shock for 45 minutes at 44°C resulted in ∼88% of L cells containing stress granules (Figure 4A). Similar to our observations in CV-1 cells pre-treated with sodium arsenite, viral protein expression was enhanced at 10 and 18 h p.i. in L cells exposed to a 45 min heat shock prior to infection (Figure 4B). However, heat shock activation had no impact on the percentage of infected cells or viral yield in L cells (Figures 4C-E). These findings suggest that in L-cells, heat shock enhances protein expression, but does not affect either the numbers of infected cells or the per cell yield of virus. Although heat shock activates HRI kinase, its effect on cells differs from sodium arsenite. Together, these data indicate that sodium arsenite pre-treatment perturbs the cell in a way that is beneficial for reovirus compared to other activators of the HRI kinase.

**Figure 4.**
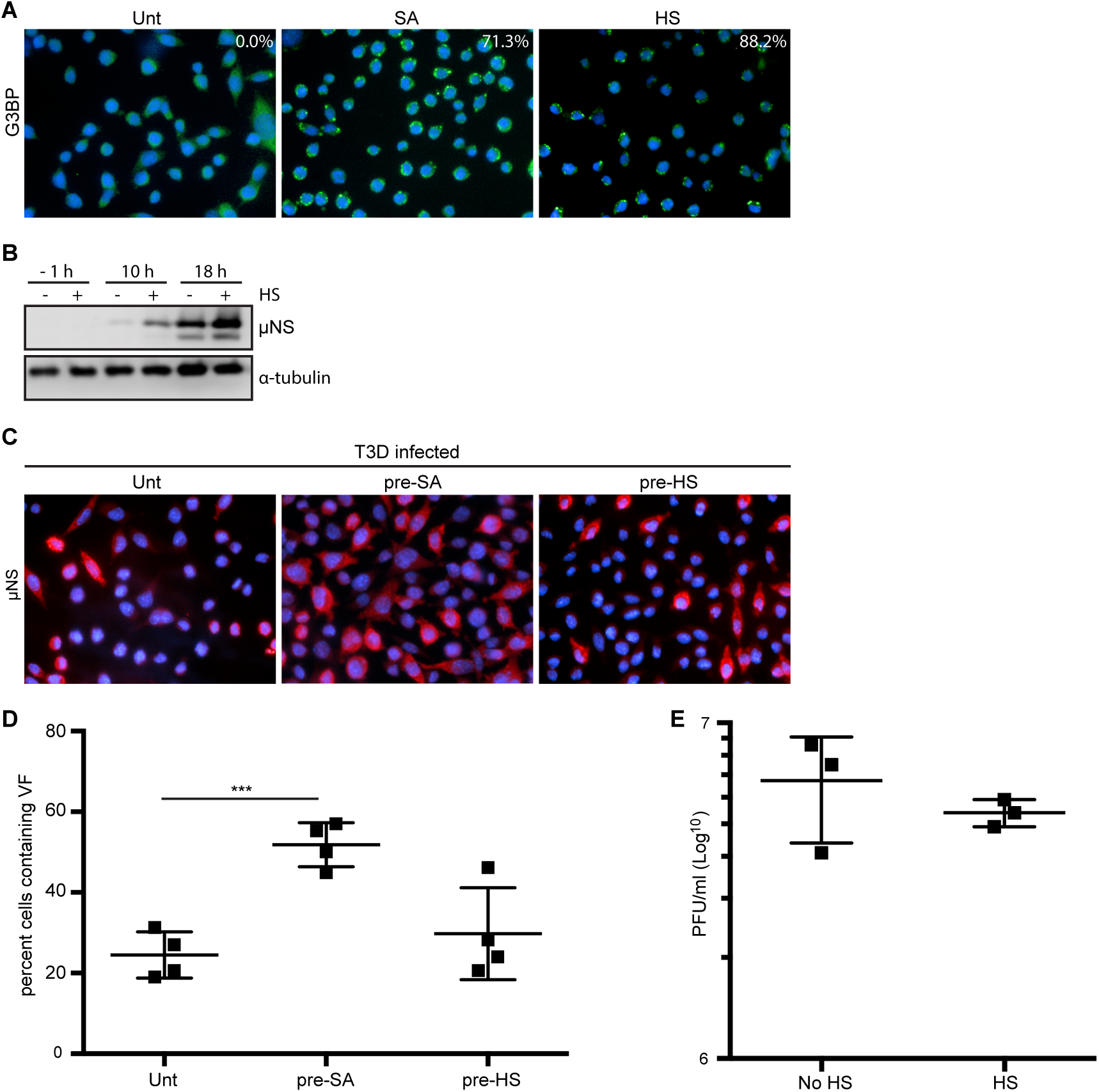
Heat shock enhances reovirus-protein expression, but not infectivity, in L929 cells. (A) L929 cells were left untreated (unt), were pre-treated with 0.5 mM SA (SA) or were heat shocked at 44°C for 45 min (HS) before fixing and immunostaining for G3BP (green) to detect stress granules (SG). The percentage of SG induced was quantified [(# of cells containing SG / total # of cells) × 100] from three independent experiments. (B) L929 cells were left untreated or were heat shocked as above immediately prior to infection with T3D at an MOI = 1. At the indicated time points, cells were lysed and the expression levels of the indicated proteins was determined by immunoblot. (C) L929 cells were left untreated, treated with 0.5 mM SA for 30 min or were heat shocked as above. Following this, cells were infected with T3D at an MOI = 1. At 18 h p.i, cells were fixed and immunostained for µNS (red) and DAPI (nuclei, blue). (D) The percent cells containing viral factories (VF) as represented in panel C was quantified [(# of cells containing VF / total # of cells) × 100] from at least three independent experiments. (E) L929 cells were left untreated or were heat shocked as above prior to infecting with T3D at an MOI = 1. At 18 h p.i., cells were subjected to the standard MRV plaque assay in L929 cells and plaques were counted from at least three independent experiments. *** *P* < 0.001; two-tailed unpaired t test.

### Addition of sodium arsenite prior to 4 hours enhances reovirus permissivity

Activation of the HRI kinase and eIF2α phosphorylation suppress general host translation [27]. We found that the enhancement of viral protein synthesis in cells pre-treated with sodium arsenite treatment was not evident if cells were treated immediately prior to harvest at 10, 18, or 24 h p.i. (Figure 2A). Given that viral protein expression, but not the percentage of infected cells or viral yield, was enhanced by activation of the HRI kinase using heat shock in L cells, we hypothesized that sodium arsenite treatment might enhance viral translation at early times post infection. To test this, CV-1 cells were either left untreated or were treated with 0.5 mM sodium arsenite at −0.5, 0, 1, 2, 4, 6, 8, or 10 h p.i. We then assessed the percentage of infected cells at 18 h p.i.. Consistent with our previous data, pre-treatment (−0.5 h p.i.) with sodium arsenite prior to inoculation with virus resulted in an increase in the percentage of infected cells (Figures 5A-B). Although pre-treatment had the greatest impact on reovirus permissivity, treatment with sodium arsenite at 0, 1, 2 and 4 h p.i. resulted in an increased percentage of infected cells as compared to untreated reovirus-infected cells (Figures 5A-B). Addition of sodium arsenite at 6 h p.i. or later had no effect on the percentage of infected cells at 18 h p.i. (Figures 5A-B). These data suggest that sodium arsenite is most beneficial when added early during the viral life cycle.

**Figure 5.**
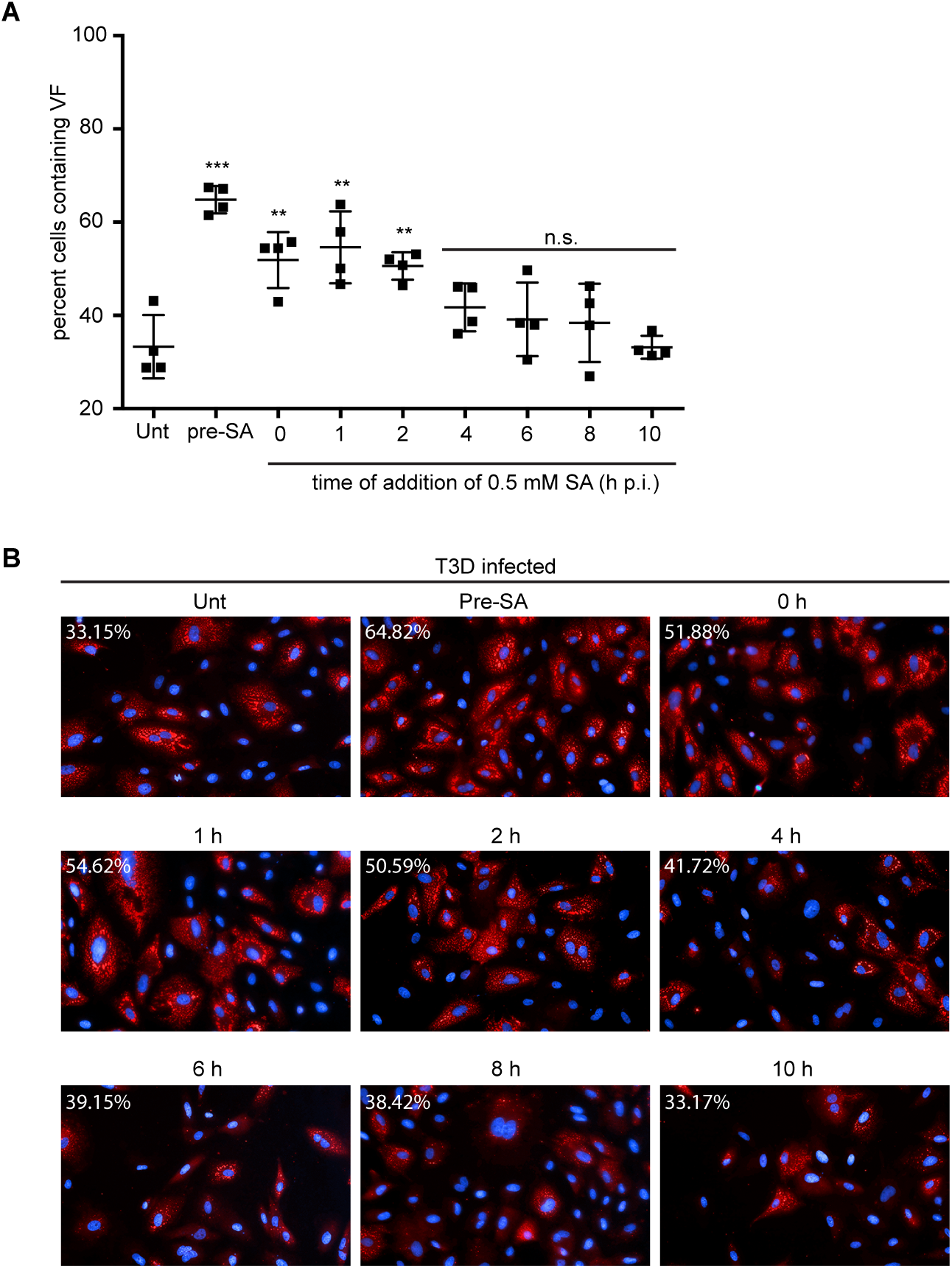
Addition of sodium arsenite prior to 4 h p.i. enhances reovirus permissivity. CV-1 cells were left untreated (Unt), were pre-treated with 0.5 mM SA for 30 min prior to virus inoculation (pre-SA) or were treated with 0.5 mM SA for 30 min at the indicated times post infection. At 18 h p.i., cells were fixed and immunostained for µNS (red) and DAPI (nuclei, blue). (B) Representative images from each of the conditions above. The percent cells containing viral factories (VF) was quantified [(# of cells containing VF / total # of cells) × 100] from four independent experiments. ** *P* < 0.01; *** *P* < 0.001; n.s. denotes not significant; two-tailed unpaired t test.

### Preventing eIF2α de-phosphorylation through the use of salubrinal has no effect on reovirus replication

Growth arrest and DNA damage protein-34 (GADD34) is expressed in response to phosphorylation of eIF2α as part of the ISR and is upregulated in cells following infection with reovirus strains that induce host shut-off [15]. GADD34 is a protein phosphatase-interacting protein that, in conjunction with protein phosphatase 1 (PP1), acts to dephosphorylate eIF2α at serine 51 during times of cellular stress [28]. Our findings thus far are consistent with previous reports suggesting that eIF2α kinase activation is beneficial to reovirus infection. We, therefore, hypothesized that reovirus may be capable of viral protein synthesis in the face of enhanced eIF2α phosphorylation. We reasoned that inhibition of GADD34/PPI dephosphorylation would enhance phosphorylation of eIF2α, but should have no effect on viral protein synthesis. We used a selective inhibitor of GADD34, salubrinal, which prevents the activity of the GADD34/PPI complex without affecting the kinases that phosphorylate eIF2α [29]. Immediately following adsorption of reovirus, CV-1 cells were treated with increasing amounts of salubrinal and cells were harvested at 18 h p.i. to determine viral protein expression (Figure 6A). Even at high concentrations, salubrinal had no impact on viral protein expression, suggesting that reovirus tolerates the cellular antiviral activity of eIF2α phosphorylation. Given that sodium arsenite is a known inducer of eIF2α phosphorylation, we also tested the impact of adding salubrinal following reovirus adsorption to sodium-arsenite pre-treated cells (Figure 6A). Again, we observed no negative impact on viral protein expression when cells were pre-treated with sodium arsenite prior to infection and then incubated in the presence of salubrinal for the duration of infection (Figure 6A). Consistent with this, the percentage of virus infected cells was unaffected by the addition of 50 mM salubrinal either in cells left untreated or cells pre-treated with 0.5 mM sodium arsenite (Figures 6B-C). Together, these data suggest that sustained eIF2α-phosphorylation resulting from salubrinal treatment during reovirus infection is not anti-viral.

**Figure 6.**
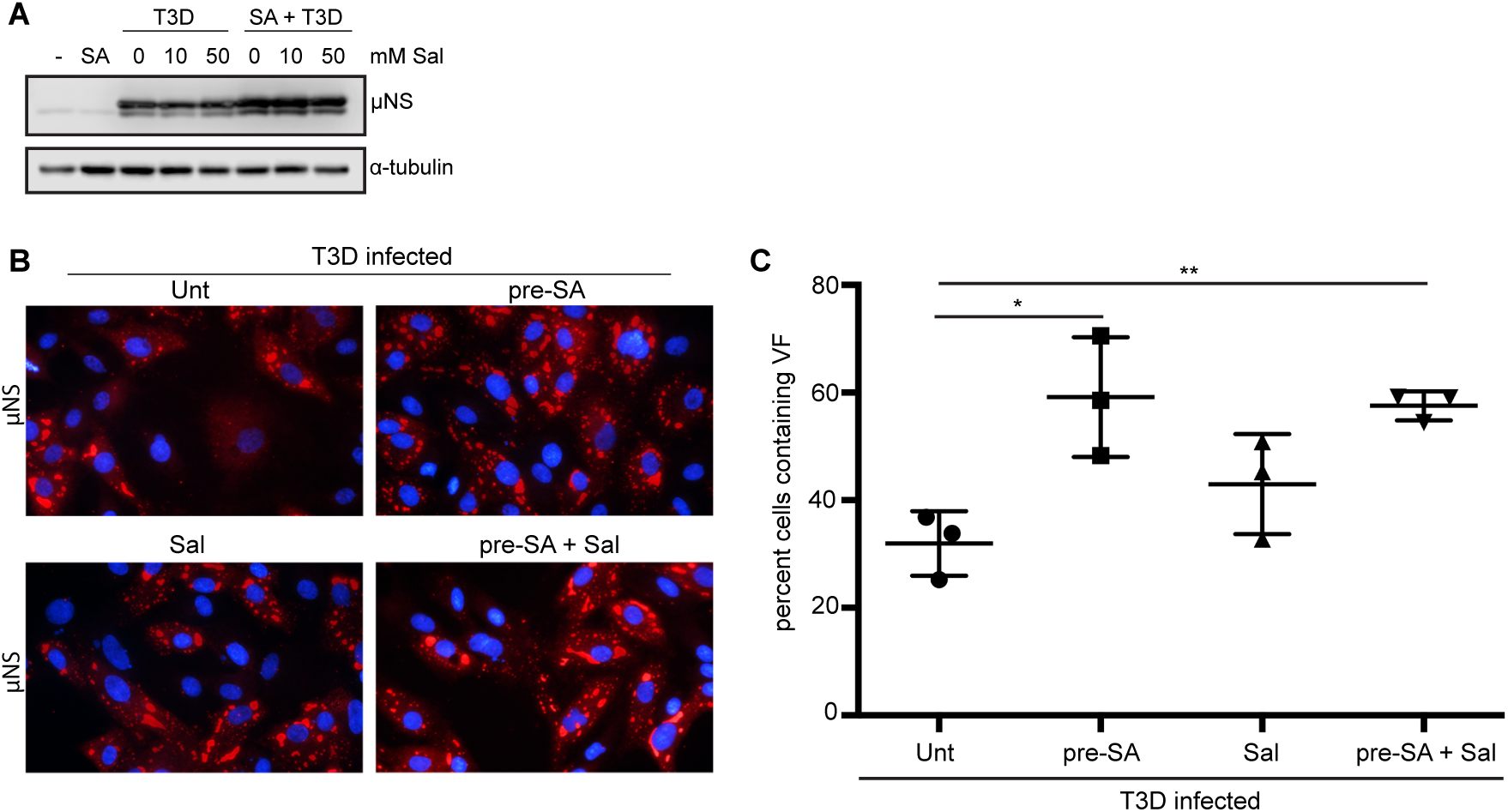
Salubrinal treatment does not negatively impact reovirus infection. (A) CV-1 cells were left untreated (-), were pre-treated with 0.5 mM SA for 30 min (SA) or were infected with T3D at an MOI = 10 in the presence or absence of SA pre-treatment. Immediately following infection, salubrinal (Sal) was added to T3D-infected cells at the indicated concentrations. At 18 h p.i., cells were lysed and the expression levels of the indicated proteins were determined by immunoblot. (B) CV-1 cells were left untreated or treated with 0.5 mM SA for 30 min prior to infection with T3D at an MOI = 1. Following infection, 50 mM salubrinal was added where indicated. At 18 h p.i., cells were fixed and immunostained for μNS (red) and DAPI (nuclei, blue). (C) The percent cells containing viral factories (VF) as represented in panel B was quantified [(# of cells containing VF / total # of cells) × 100] from 3 independent experiments. * *P* < 0.05, ** *P* < 0.01; two-tailed unpaired t test.

## DISCUSSION

In this report, we find that treatment of cells with sodium arsenite (SA), a potent inducer of the integrated stress response (ISR) and eIF2α phosphorylation, prior to inoculation with reovirus is beneficial to the virus in a strain-independent but cell-type-specific manner. SA treatment induces the formation of reactive oxygen species (ROS) within cells. The increased intracellular ROS in turn lead to activation of HRI, which then phosphorylates eIF2α at serine residues 48 and 51 leading to general translation repression and downstream activation of the integrated stress response. In order to mature and become activated, HRI has to be in a complex with CDC37, PPP5C, and HSP90. In addition to being activated by heme-deficiency, HRI is also activated by heat shock and osmotic shock, but not by ER stress or by amino acid or serum starvation [7]. In contrast, to the enhanced viral replication we saw following pre-treatment with SA, heat shock (HS) pre-treatment did not enhance viral replication. Heat shock also activates HRI, but to a qualitatively lower level than SA and with a different pattern of autophosphorylation of HRI [7]. Why pre-stressing cells with SA, but not with heat shock, benefited viral replication is unclear. It is possible that the kinetics, magnitude and timing of eIF2α phosphorylation following these different stressors varied leading to the different outcomes we saw. Alternatively, we cannot rule out the possibility that SA treatment activates responses not mediated by HRI that benefit viral replication.

In addition to inducing eIF2α phosphorylation, SA induces the formation of stress granules (SG). Consistent with the findings of Qin et al, we found that reovirus infection induced SG formation early after infection with the number of SG reducing by 8 h and disappearing late in infection. Our data in CV-1 cells are not consistent with the report by Smith et al, who detected SGs at 19.5 h post-infection of DU145 cells [15, 16]. While it is likely that different strains of reovirus induce SG to varying extents, and that this is influenced by cell type and MOI, our data using the type 3 Dearing strain suggests that infection with this strain of reovirus results in a low level of SG induction that is absent at late times post infection. The timing of SG appearance (peak at 6 h) and dissolution (beginning at 8 h) in our hands supports the idea put forth by Qin et al that SG induction is linked to VF formation and that the onset of viral translation interferes with SG stability [16]. Smith et al stained reovirus-infected cells at 19.5 h post-infection for the SG protein, TIAR, but did not co-stain for a viral protein, such as µNS, to detect viral factories. We and others have observed that the SG protein G3BP1 localizes to the outer margins of viral factories at late times post infection in a fraction of reovirus infected cells [17]. Therefore, the TIAR-positive punctae seen at 19.5 h post-infection by Smith may in fact have been viral factories that were co-staining for the presence of TIAR [15]. The significance of SG protein relocalization to VFs remains to be determined. Still, **t**hese data imply that reovirus has evolved mechanisms to counter the cellular antiviral activity of translation suppression through stress granule induction.

Pre-treatment of cells with SA led to enhanced viral protein expression, increased infectivity, and higher viral titers. Given that reovirus compartmentalizes the translational machinery in VF and that components of the translational machinery are also sequestered within SGs, we previously speculated that SGs may serve as a reservoir of translational machinery for reovirus VFs [23]. However, comparison of reovirus infection following pre-treatment with either SA or heat shock (HS) suggests that the benefit is independent of SG formation, as both treatments resulted in a robust induction of SGs, but only SA pre-treatment led to an increase in viral replication. As we only examined viral replication late during infection, it is possible that we missed early differences. Additionally, since we were unable to achieve robust SG formation following HS in CV-1 cells, viral replication studies were performed in L cells. Therefore, we can’t rule out cell-specific influences.

Our data is consistent with the findings of Smith et al that reovirus benefits from activation of the ISR. In that study, the authors found that reovirus replicated to lower titers in cells lacking a phosphorylatable eIF2α [15]. SA is a potent activator of the ISR. SA-mediated activation of the HRI kinase leads to phosphorylation of eIF2α and downstream upregulation of ATF4 and other transcriptional mediators of the ISR [27]. Given that canonical cap-dependent cellular translation relies on the availability of active GTP.eIF2, it is possible that activation of the ISR allows reovirus mRNAs a competitive advantage for limited translational machinery. The effect of SA was greatest when added before reovirus adsorption but was still beneficial if treatment occurred within the first 4 h of infection. Translation of reovirus mRNAs rises sharply at ∼ 6 h p.i. and peaks at ∼12 h p.i. [30]. Given that the rising level of viral protein synthesis is concomitant with continued host protein synthesis, it is possible that SA treatment selectively reduces the translational machinery available for host protein synthesis [30]. It has been postulated that heightened eIF2α-phosphorylation may more strongly affect mRNAs containing long or highly structured 5’ untranslated regions (UTRs) [31, 32]. Reovirus mRNAs have short 5 ′ and 3 ′ UTRs, possibly providing protection to viral mRNAs during translation [33, 34]. Our findings that reovirus protein expression and infectivity were unaffected when we treated cells with salubrinal to prevent de-phosphorylation of eIF2α further supports this and suggests that reovirus is refractory to salubrinal exposure.

It is known that under times of stress and reduced availability of active eIF2, cells can utilize other initiation factors to ensure translation of select mRNAs. These factors, including eIF2A, eIF2D, ligatin, MCT-1/DENR, and N-terminally truncated eIF5B_479-1220_, can promote efficient recruitment of Met-tRNA^Met^i to the 40S/mRNA complex under varying circumstances [35-39]. Several viruses take advantage of these alternative initiation pathways including Sindbis virus and poliovirus [36, 39]. We have noted that two of these factors, eIF2A and eIF5B localize to viral factories within infected cells (unpublished observations, JSLP). As yet, we do not know if these factors play a functional role in translation of viral mRNAs within VFs or if they are being sequestered to prevent cellular mRNA translation.

We cannot exclude the possibility that pre-treatment of cells with SA has other effects that promote viral replication independently of eIF2α phosphorylation. Reovirus is an oncolytic virus, preferentially infecting cancer cells. This phenomenon has been linked to a mutationally-active Ras pathway, however reports have been conflicting regarding Ras dependency [40-43]. Cancer cells have elevated levels of chaperones that facilitate the high levels of protein synthesis typical of transformed cells. The efficiency of mRNA translation is increased in the presence of supplemental recombinant Hsc70 and the rate of translation elongation may be regulated by chaperone availability [44]. Treatment with SA also increases Hsc70/Hsp70 chaperone levels in cells [45]. Furthermore, Hsc70 is required for HRI activation and blockade of Hsc70 disrupts HRI activation [7]. We have previously reported that Hsc70 is specifically targeted to the viral factory, although why this protein is specifically recruited remains unclear as its recruitment is independent of its chaperone function [46]. It is conceivable that SA pre-treatment could alter the availability of Hsc70 or other protein folding chaperones in a way that is favorable during reovirus replication.

Infection with reovirus results in increased expression of stress response genes including Hsc70, and GADD34, the latter of which complexes with protein phosphatase 1 to reverse eIF2α phosphorylation. We observed that Type 3 Dearing (T3D) reovirus protein expression was unaffected in the presence of salubrinal, a selective inhibitor of the GADD34/PP1α complex responsible for dephosphorylating eIF2α, even in cells pre-treated with SA to elevate the level of phosphorylated eIF2α. The observation that salubrinal was not anti-viral to T3D reovirus is not unique. Rotavirus, a fellow member of the Reoviride family, as well as other viruses including hepatitis C virus and mouse hepatitis coronavirus, demonstrate continued viral protein synthesis despite heightened eIF2α-phosphorylation [47-49]. While this provides additional evidence that reovirus benefits from the ISR, it is important to note that reovirus-induced expression of stress response genes is not uniform and that host shutoff strains, including C8 and C87 (Type 3 Abney, T3A), induce increased expression of these genes compared to T3D, which is not considered a host shutoff strain. While all reovirus strains tested, including T3A, benefited from SA-pre-treatment, host shutoff strains may respond differently from T3D to salubrinal treatment [15].

In addition to strain-specific differences in the ability to induce host shutoff, cell differences have also been recorded. For instance, reovirus-induced host shutoff was not observed in HeLa cells but was in L cells [30]. This may be linked to our observation that T3D infection in HeLa cells is unaffected by SA pre-treatment, whereas SA pre-treatment is beneficial in other cell types including L cells, CV-1 and HPDE cells.

Overall, this study finds that sodium arsenite-induced activation of the ISR prior to reovirus inoculation results in enhanced viral protein expression, increased infectivity and higher viral yield. Furthermore, HRI kinase activation of the ISR is insufficient for replication enhancement as both heat shock and sodium arsenite activate the HRI kinase, but only sodium arsenite treatment was associated with increased viral infectivity and higher yield. Finally, sodium arsenite-induced enhancement was observed across reovirus strains but was not observed in all cell types suggesting a role for cell-type-specific influences. Understanding the interplay between reovirus and the ISR, and how to modulate it to enhance virus replication, could reveal novel targets to strengthen the oncolytic potential of reovirus.

## ACKNOWLEDGEMENTS

Work in E.D.L.’s lab was supported by Le Moyne College Research and Development Grants (to E.D.L), and Le Moyne College Student Research Grants awarded to M.M.L and M.P.W..

