## Supplemental Figure 1 for "*Mammalian orthoreovirus* infection is enhanced in cells pre-treated with sodium arsenite"

**Supplemental Figure 1. Pre-treatment with 0.5 mM sodium arsenite enhances reovirus permissivityin L929 cells but not HeLa cells across reovirus.** (A) L929 or (B) HeLa cells were left untreated (No SA) or were treated with 0.5 mM SA for 30 min prior to infection (Pre-SA). Following this, cells were reovirus strains , T1L or T3A such that ~20% of cells were infected. At 18 h p.i., cells were fixed and immunostained for μNS (red) and DAPI (nuclei, blue). The percentcells containing viral factories (VF) was quantified [(# of cells containing VF / total # of cells) × 100]. * *P* < 0.05; ** *P* < 0.01; two-tailed unpaired t test.

**SUPPLEMENTAL FIGURE 1**

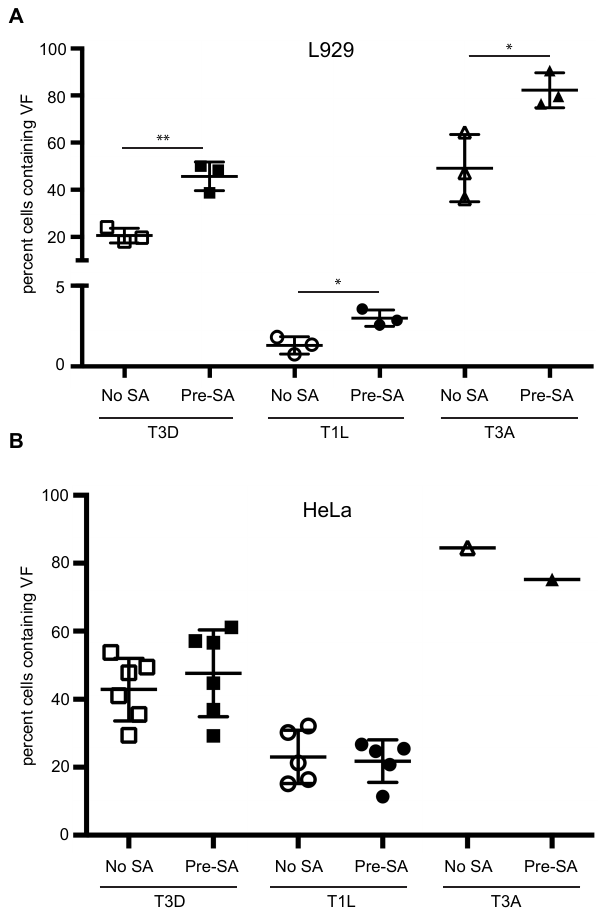
